# Subtropical anticyclones shape life-history traits of a wind-reliant marine top predator

**DOI:** 10.1101/2025.02.27.640559

**Authors:** Ruijiao Sun, Etienne Rouby, Christophe Barbraud, Henri Weimerskirch, Karine Delord, Kristen Krumhardt, Caroline C. Ummenhofer, Stéphanie Jenouvrier

## Abstract

Subtropical anticyclones are semi-permanent atmospheric high-pressure systems located in all major ocean basins and are associated with large-scale wind and weather patterns. They shape the physical environments of many species, yet their demographic impacts on wild populations remain largely unexplored. We combined population and climate analyses to investigate the demographic effects of subtropical anticyclones. Using 39 years of monitoring data on wandering albatrosses (*Diomedea exulans*), the “wind rider”, breeding in the southern Indian Ocean, we explored the mechanisms linking subtropical anticyclone with demographic rates. We found that an intensified and poleward-shifted Indian Ocean subtropical anticyclone enhances westerly winds, improving survival and reproduction across all life stages in wandering albatrosses. These findings uncover a direct link between subtropical anticyclones and population dynamics, highlighting subtropical anticyclones as potential drivers of wind-reliant taxa responses to climate variability and change.

## 1 Introduction

Subtropical anticyclones are semi-permanent high-pressure systems that span approximately 40% of the Earth’s surface, centered around 30^◦^ latitude in both the Northern and Southern Hemispheres (Rodwell & Hoskins, 2001). They link the mid-latitude westerly wind regime and tropical easterly trade winds through anticyclonic atmospheric circulation (Miyasaka & Nakamura, 2010), and are associated with lateral and vertical gradients in atmospheric structures that shape the global climate (Rodwell & Hoskins, 2001). By driving large-scale weather systems, subtropical anticyclones have been shown to influence species behaviors, such as migration patterns of the brown planthopper (*Nilaparvata lugens*) (Hu et al., 2025; Zhang et al., 2025), and may ultimately affect population dynamics. Although variability in subtropical anticyclones alters winds, oceanographic conditions, and food availability, there is limited empirical evidence directly linking these systems to survival and reproduction in wild populations.

Variability in the location and intensity of subtropical anticyclones drives changes in wind direction and speed, which influence the movement and energy expenditure of many species. These wind-driven changes can affect demographic rates in species that respond behaviorally or physiologically to wind variability, such as by adjusting foraging strategies to improve efficiency and conserve energy (Hennessy et al., 2020; Thorne et al., 2023; Wijers et al., 2022). Furthermore, winds can influence predator-prey interactions by shifting prey distribution (Gangoso et al., 2020). Subtropical anticyclones could also shape the demographic rates of wild populations through bottom-up mechanisms. Indeed, changes in wind associated with subtropical anticyclones influence the intensity of oceanic Ekman upwelling, impacting primary production and resource aggregation (Bograd et al., 2023; Rykaczewski et al., 2015). Mid-latitude weather systems in both hemispheres are closely linked to anticyclone variability through its influence on atmospheric blocking, regional winds (Stowasser et al., 2007), SST (Chung et al., 2011), and precipitation (Li et al., 2011), all of which impact large-scale primary production and drive bottom-up effects.

Subtropical anticyclones can influence a wide range of taxa. Winds on the west coast of North America are primarily affected by the North Pacific High, and have been shown to impact the survival of Nearctic-neotropical migrating yellow warbler (*Setophaga petechia*) (Drake et al., 2014). Variability of the North Pacific High also drives biological responses in the California Current system (Schroeder et al., 2013), and winds have been linked to the distribution and abundance of blue whales (*Balaenoptera musculus*) through bottom-up effects (Ryan et al., 2022). The Azores High (or Bermuda High) and South Pacific High are also linked to highly productive eastern boundary upwelling systems, namely the Canary Current and Humboldt Current systems (Barton et al., 1998; Mogollón & R. Calil, 2018). Thus, subtropical anticyclones can affect population dynamics through a combination of winds and bottom-up mechanisms. This highlights the complex interplay between biology and physical processes, emphasizing the importance of studying the impacts of subtropical anticyclones across taxa and regions.

Many subtropical anticyclone systems are associated with large-scale climate mode indices. For instance, the variability of the North Atlantic Oscillation (NAO), an atmospheric dipole, is represented by the difference in sea-level pressure between the Azores High and the Icelandic Low (Hurrell & van Loon, 1997). However, the expansion of the Azores High has been strongly associated with increased drought in the Iberian Peninsula, whereas the NAO has shown weaker and less consistent links to regional hydroclimate (Cresswell-Clay et al., 2022). Similarly, the Western Pacific Subtropical High is a robust predictor of monsoon strength and tropical storm activity (Wang et al., 2013; Yang et al., 2022). Therefore, focusing on subtropical anticyclone systems independently offers a subtropical perspective that provides a more direct and precise understanding of regional climate variability.

The intensity and extent of subtropical anticyclones encapsulate environmental variability in the coupled ocean-atmosphere system that is challenging to link directly to demographic patterns. This aligns with the advantage of using large-scale climate indices, which aggregate environmental conditions and have been used extensively to provide a comprehensive understanding of ecological and demographic responses to environmental variability (Stenseth et al., 2003). Unlike widely studied climate modes, such as the NAO and El Niño–Southern Oscillation (ENSO), which have readily available predefined indices (Hurrell et al., 2003; McPhaden et al., 2006), subtropical highs lack standardized indices (but see the use of Western Pacific Subtropical High indices in Lv et al. (2023)). Instead, climatologists often develop custom approaches to describe subtropical anticyclones based on specific research objectives (Miyasaka & Nakamura, 2010; Rodwell & Hoskins, 2001). The limited investigation of subtropical anticyclones in ecology likely arises from the perceived complexity of quantifying the variability of these high-pressure systems.

Here, we use indices describing the variability in the strength and location of a subtropical anticyclone system, and test its impacts on the life-history traits of a marine top predator. We utilize a 39-year (1980-2018) capture-mark-recapture dataset of a wandering albatross (*Diomedea exulans*) population breeding in the southern Indian Ocean. Climate and oceanographic conditions in this region including wind, sea surface temperature (SST), and precipitation are strongly influenced by the Mascarene High (MH), the South Indian Ocean subtropical anticyclone (Manatsa et al., 2014; Morioka et al., 2015). The wandering albatross is one of the most wide-ranging pelagic seabirds, known for utilizing wind to undertake long-distance foraging trips (Richardson, 2011), with strong winds enhancing their foraging efficiency (Cornioley et al., 2016; Weimerskirch et al., 2012). Their diet consists primarily of cephalopods (Xavier et al., 2014), which have fast life-history traits and show rapid biomass responses to oceanographic conditions and primary production within approximately six months (Arkhipkin, 2004; Hoving et al., 2013; Marcout et al., 2024). An even faster response may occur in the small-sized squid targeted by albatrosses. Previous research has suggested that wind conditions influence foraging efficiency and breeding success, but has struggled to identify a suitable wind index to quantify correlations with demographic rates likely due to spatial and temporal mismatches, as albatrosses forage across vast areas (Weimerskirch, 2018; Weimerskirch et al., 2012). Employing subtropical anticyclone indices, which encapsulate large-scale atmospheric and oceanographic conditions, can offer new insights into how highly mobile species respond to climate change.

We apply Bayesian multi-event capture–mark–recapture models to quantify the effects of MH on the full life cycle of wandering albatrosses, including adult survival, breeding probability, and breeding success, as well as the probability of first breeding attempt and first breeding success in juveniles. We also examine the effects of ENSO, which has been shown to influence the biology of this wandering albatross population (Barbraud et al., 2012), and the Indian Ocean Dipole (IOD) index, which affects large-scale climate in the southern Indian Ocean (Saji & Yamagata, 2003). Using climate anomaly composite analyses, we identify the mechanisms by which the subtropical anticyclone system shapes the life-history traits of this marine top predator. Oceanographic variables were derived from observation-based climate reanalysis datasets, alongside a model-based reconstruction of ocean primary productivity.

## 2 Materials and methods

### 2.1 Study system and data collection

Wandering albatrosses forage in subantarctic and subtropical waters between latitudes 30^◦^ S and 60^◦^ S while breeding (Weimerskirch et al., 2012), and show a wide distribution in the Southern Ocean during the nonbreeding period (Weimerskirch et al., 2005), mainly feeding on fish and squid. Our study population of wandering albatrosses breed on Possession Island, Crozet (46^◦^24’S 51^◦^46’E) where a capture-mark-recapture (CMR) program has been undertaken since 1965. Fledglings and adults are banded annually using individually coded stainless-steel leg rings. Reproductive status was determined during the breeding season. Between January and February after egg laying, nests are visited three to four times to identify breeding individuals. Chicks are ringed in September and October before fledging. Breeding success is determined based on chick survival until fledging. Unmarked individuals found at the colony are also ringed each year. Sex is identified based on morphology and genetic assessment (Weimerskirch et al., 2005). In this study, we relied on data collected from 1980 to 2018, totaling 11584 individuals.

### 2.2 Mascarene High indices

The Mascarene High (MH), located in the southern Indian Ocean, is a semi-permanent subtropical anticyclone system with the potential to influence weather and climate over broader regions. Therefore, in this study, we examined how variations in MH characteristics impact the vital rates of our study species, focusing on changes in MH intensity and location (latitude and longitude). We utilized monthly reanalyzed sea level pressure data from ERA5 to calculate MH indices (Hersbach et al., 2020). The MH area was defined as the total area within the region 30^◦^E–120^◦^E and 20^◦^S–50^◦^S above a sea level pressure of 1020.0 millibars contour (Appendix Fig. S1). The center of the MH is identified as the location with the highest sea-level pressure within the MH area. The MH strength is defined as the average sea level pressure value within the 1020.0 millibars contour, and the latitude and longitude of the MH center represent the spatial location of the MH (Appendix Fig. S1).

### 2.3 Bayesian multi-state capture-mark-recapture model (MSCMR)

We used the same Bayesian MSCMR model as in Van de Walle et al. (2024) to estimate survival, breeding probability, and breeding success probability of wandering albatrosses. In a life cycle of wandering albatrosses, individuals move between a set of states over a time step of a year, conditional on transition probabilities. The wandering albatross states depend on age classes and reproductive status. All vital rates vary with age and states and we followed the definitions in Van de Walle et al. (2024). We used the same Bayesian MSCMR model as in Van de Walle et al. (2024) to estimate survival, breeding probability, and breeding success probability of wandering albatrosses.

We constructed and ran separate models for female and male life histories to maintain statistical independence. Although the full dataset spans 1965–2018, we focused our analysis on the period from 1980 to 2018 to assess the impact of the MH on demographic rates. This period corresponds to improved banding and more robust monitoring, yielding higher-quality population data. We also relied on ERA5 reanalysis data to calculate MH indices and extract oceanographic conditions and the inclusion of satellite observations starting in 1979 enhances the reliability of ERA5 from that point onward (Hersbach et al., 2020). To improve the precision of vital rate estimates, we leveraged the full demographic dataset and modeled juvenile and adult components separately. We modeled the juvenile and adult components of the population separately to estimate their respective demographic rates. Individuals born after 2016 were excluded from juvenile analyses due to low detection rates between ages 1 and 5, which could lead to underestimates of survival. To reduce model complexity and computation time, we tested the effects of MH indices on each demographic rate independently, resulting in 36 total models: 9 models for survival, breeding probability, and breeding success in adults of each sex, and 9 for the same rates in juveniles of each sex.

To evaluate MH effects on vital rates, we defined season-specific covariate windows: MH conditions averaged from January to December for survival, September to December (pre-laying period) for breeding probability, and January to October (incubation to chick-rearing) for breeding success. We modeled the MH indices as additive temporal covariates. All covariates were detrended and normalized prior to analysis.

Multistate capture–mark–recapture (MSCMR) analyses for adult individuals were conducted in JAGS (Plummer, 2003) via R (R Core Team, 2021) using the jagsUI package (Kellner & Meredith, 2021), and analyses for juveniles were performed using NIMBLE (de Valpine et al., 2017). Key parameter estimates were compared across platforms to ensure consistency. We used the R Package “nimble” version 1.01 on R version 4.3.1. For each model, we ran 3 parallel chains with 20,000 iterations, a burn-in phase of 4000, and a thinning interval of 2 for a total of 24,000 iterations. Convergence was confirmed by visual examination of the posterior distributions and the Gelman-Rubin statistic, with a R-hat lower than 1.05 indicating that convergence was reached (Brooks & Gelman, 1998). The confidence of an effect was calculated as the proportion of the posterior distribution that shared the same sign as the posterior mean (Jeffreys, 1998; Ly et al., 2016), which we referred to as the ‘f’ value (Fay et al., 2022; Van de Walle et al., 2024). An f-value near 100% signifies strong evidence of an effect, whereas values around 50% indicate no clear evidence. Here, we categorized effects as significant using a 90% threshold.

### 2.4 Climate anomaly composite analysis

We used monthly reanalyzed SST, sea level pressure, and surface wind data from ERA5 (Hersbach et al., 2020). Ekman upwelling was calculated following Kessler (2002) using surface wind data. Net primary production (NPP) data were obtained from the output of the Community Earth System Model version 2 (CESM2) (Danabasoglu et al., 2020; Danabasoglu, 2019), which includes an ocean biogeochemistry model (Long et al., 2021), and was forced by the Japanese 55-year Reanalysis (JRA-55) (Kobayashi et al., 2015). CESM, run in an ocean–sea-ice-only configuration, can effectively reconstruct past ocean and sea-ice conditions when these components are forced by historical atmospheric observations (i.e., atmospheric reanalysis data like JRA-55). In addition to physical ocean variables, ocean biogeochemistry and lower trophic level ecosystem dynamics are simulated within the ocean component, the Marine Biogeochemistry Library (MARBL) (Long et al., 2021). This provides variables such as NPP, which often lack complete observational records, especially in remote regions like the Southern Ocean. We used the vertically integrated total carbon fixation between 0 and 100 m depth as a measure of NPP.

We used composite anomaly analysis to investigate the mechanisms by which the intensity and location of the MH affect the vital rates of wandering albatrosses. Long-term mean climate conditions were computed as a reference baseline. We then identified years corresponding to the ten highest and ten lowest values (i.e., upper and lower 25th percentiles) of each vital rate of interest, including survival, breeding probability, and breeding success. For each vital rate, low anomaly composites were calculated by averaging conditions during the ten years with the lowest values and subtracting the overall mean conditions. Similarly, high anomaly composites were calculated by averaging conditions during the ten years with the highest values and subtracting the overall mean conditions. These composites represent the anomalous climatic conditions during years with high or low vital rates, relative to the long-term mean conditions. Significance was assessed using bootstrapping. Specifically, ten years of data were randomly drawn 1,000 times, and mean conditions were subtracted to create a distribution. Statistical significance was determined based on the 90th percentile of this distribution. The significance of the difference between high and low anomaly composites was evaluated by randomly dividing the data into two groups of 10 years, subtracting each other, and repeating this process 1,000 times. The 90th percentile of the resulting distribution was used to determine significance.

## 3 Results

### 3.1 Mascarene High affects survival and reproduction of wandering albatross

We examined how variations in the strength and spatial location of the MH (Appendix Fig. S1) impact the survival, breeding, and breeding success probabilities of wandering albatrosses breeding in the southern Indian Ocean (Fig. 1). We found that the effects of MH on wandering albatrosses varied by sex and life stage (Fig. 1). Both the strength and spatial location of the MH significantly influenced the vital rates of wandering albatrosses (Appendix Table S1 and S2 for full model results). An equatorward shift of the MH was associated with decreased vital rates, including a decrease in adult male survival probability (slope = −0.132, confidence f-value [f] = 97.0%), breeding probability for both females (−0.131, f = 98.3%) and males (−0.102, f = 98.1%), and juvenile male first-time breeding success (−0.102, f = 92.7%) (Fig. 1A). On the other hand, an intensified MH was positively correlated with adult male survival (0.129, f=95.7%), breeding probability (0.105, f=97.5%), and juvenile female first-time breeding success (0.117, f=90.8%) (Fig. 1B). An eastward shift of MH was positively correlated with increased juvenile male survival (0.205, f=95.7%) and first-time breeding success (0.135, f=96.6%) probabilities (Fig. 1C). Additionally, positive phases of ENSO and IOD were correlated with higher adult male survival, and negative phases of the IOD were linked to reduced juvenile female survival (Appendix Table S1).

**Figure 1.**
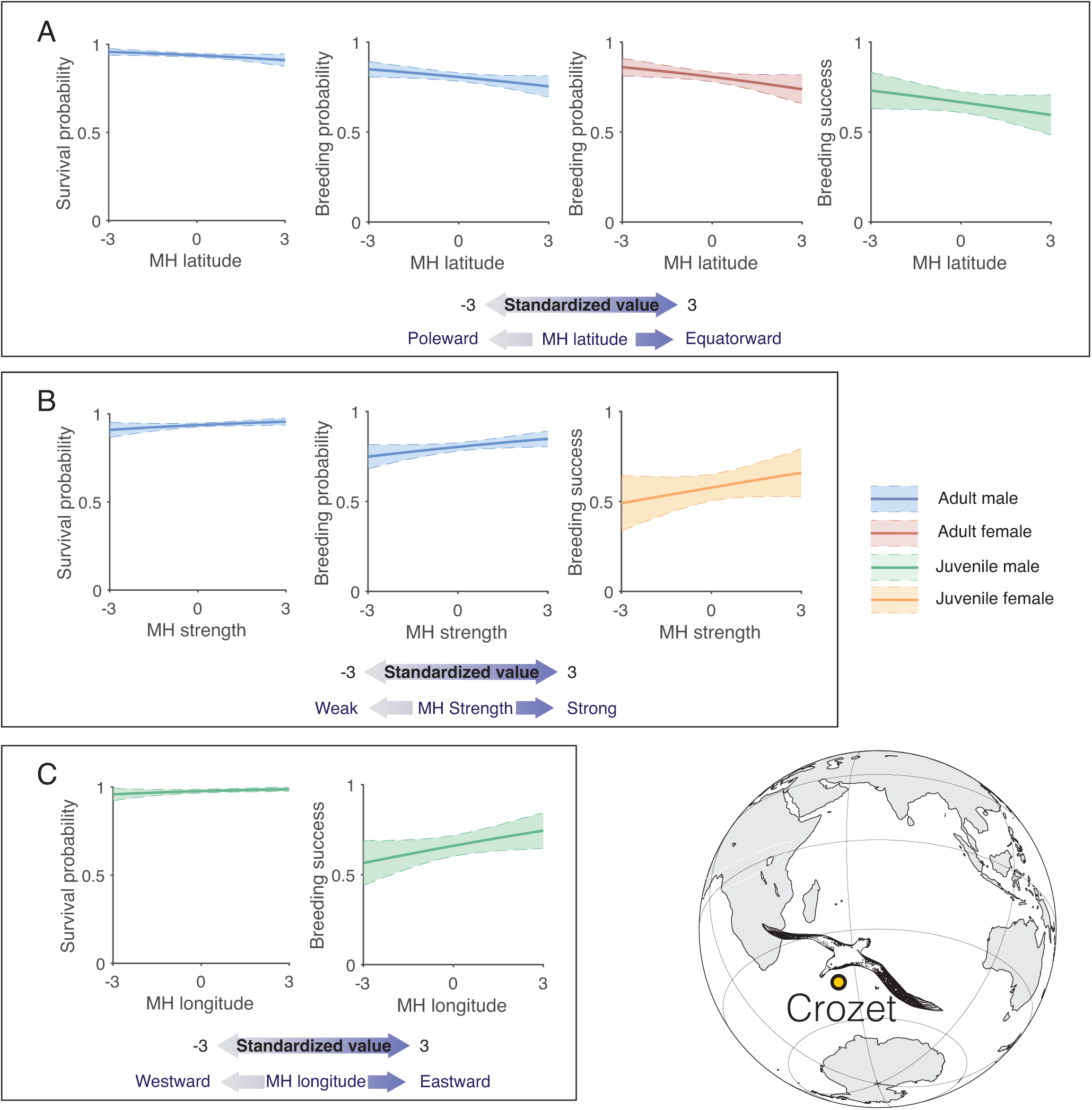
Effects of variability in Mascarene High (MH) intensity and spatial location on the life-history traits (*±*95%) of wandering albatrosses breeding on Crozet Island. (A) Effects of MH latitude on vital rates; (B) Effects of MH strength on vital rates; (C) Effects of MH longitude on vital rates. Note that the breeding success probability for juveniles refers to breeding success of their first breeding attempt in life. All covariates have been standardized. An increase in MH latitude indicates an equatorward shift of the center, while increased MH strength signifies an intensified MH, and increased MH longitude indicates an eastward shift of the MH center.

Adult males were significantly influenced by the variability of the MH in terms of both breeding probability and survival, while it only affected the breeding probability of adult females. Furthermore, when comparing juveniles to adults, the MH impacted adult breeding probability without affecting breeding success, whereas it influenced juvenile breeding success without impacting breeding probability.

### 3.2 Mechanisms linking Mascarene High variability to demographic rates of wandering albatross

To investigate the underlying mechanisms, we performed composite anomaly analyzes comparing environmental conditions during years with high versus low values of vital rates (see Methods Section 2.4 for details). These analyzes revealed associated patterns in sea level pressure, winds, SST, upwelling, and primary production, providing insight into why some years result in higher vital rates while others correspond to lower demographic performance. We also performed similar composite anomaly analyzes for MH index time series to establish their links to wind, SST, upwelling, and primary production (Appendix Figs. S7–S9). We found that an intensified and poleward-shifted MH strengthens westerly winds and increases overall wind stress across the southern Indian Ocean (Appendix Figs. S7A and S8A), whereas longitudinal shifts in the MH primarily modify the meridional wind distribution (Appendix Fig. S9A).

#### 3.2.1 Strength and location of the Mascarene High alter large-scale wind conditions

We first investigated sea level pressure and wind conditions. Here we present only the anomaly composite analysis for adult male survival (Fig. 2), while all other results, which demonstrated similar and consistent patterns, are included in the Appendix (Fig. S2-6). Years with lower vital rate values were associated with low sea level pressure anomalies in both the subtropical and subantarctic zones of the southern Indian Ocean (Fig. 2B), indicating a weakened, equatorward, and westward shift of MH. The low sea level pressure anomaly resulted in a weaker meridional pressure gradient, leading to reduced westerly winds (Fig. 2B). Conversely, years with higher vital rates were associated with high sea level pressure anomalies in both the subtropical and subantarctic zones, along with low anomalies around 60-70^◦^S (Fig. 2C). This increased the meridional pressure gradient in the subantarctic zone and enhanced westerly winds (Fig. 2C). Overall, an intensified, poleward, and eastward shift of the MH, enhancing the westerly wind belt through an increased meridional pressure gradient, was associated with higher survival, breeding, and breeding success probabilities (Fig. 2D, Appendix Fig. S2-6).

**Figure 2.**
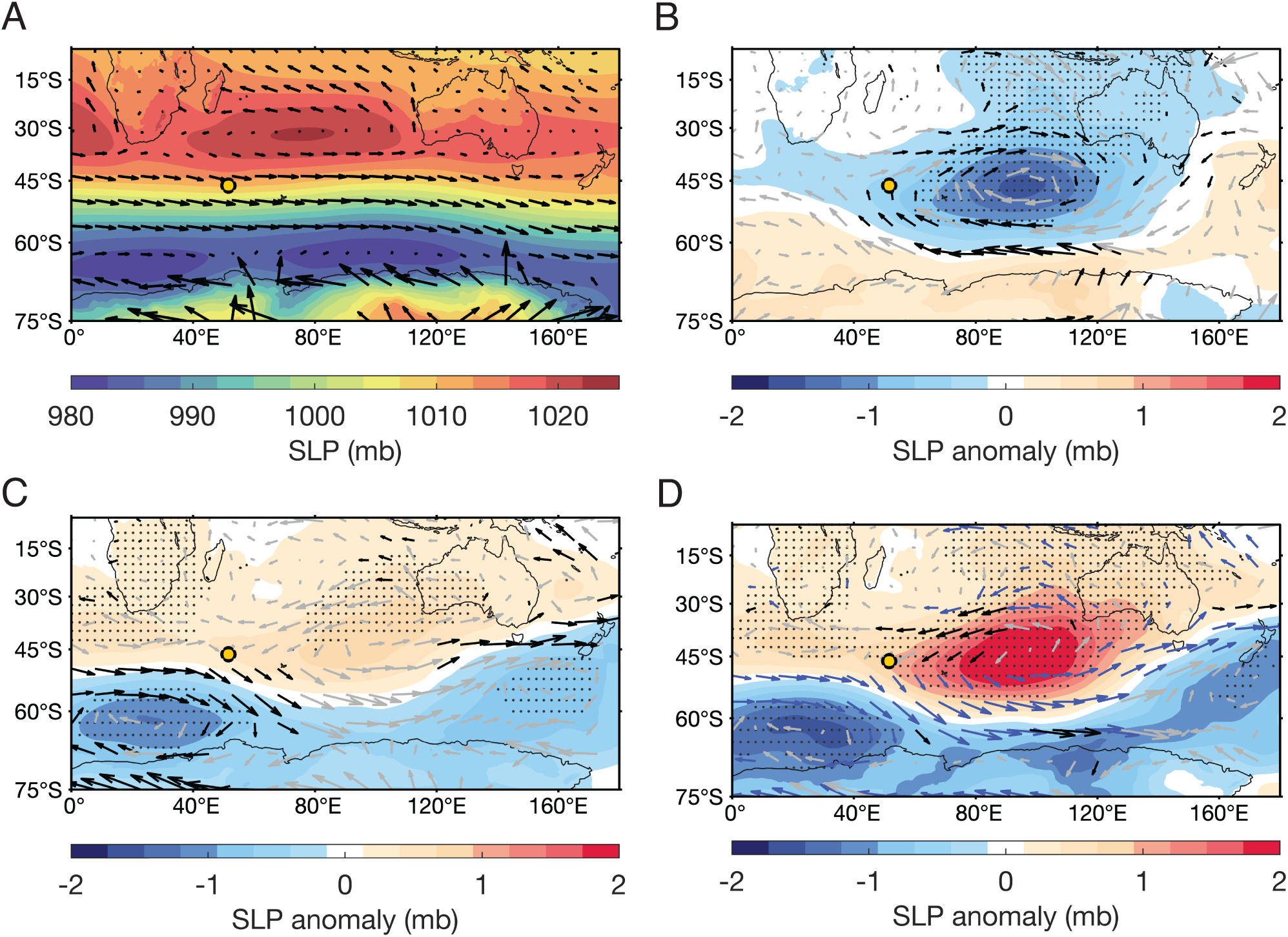
Composite anomaly analysis of sea level pressure (SLP) and wind conditions affecting the survival probability of male wandering albatrosses. (A) Long-term mean (1980-2018) of SLP and wind conditions. (B) Composite anomalies of SLP and winds during years with low male survival probability (lower 25th percentile), and (C) years with high male survival probability (higher 25th percentile). Black wind vectors indicate significant wind anomaly and light gray wind vectors indicate insignificant wind anomaly. Light dotted stippling indicates significant anomaly in SLP. (D) Differences in SLP and wind conditions during composites of high and low male survival years. Black wind vectors indicate significant wind differences associated with reduced wind speed, blue wind vectors indicate significant differences with increased wind speed, and light gray wind vectors indicate insignificant differences. Light dotted stippling denotes significant differences in SLP.

#### 3.2.2 Bottom-up effects

We investigated potential bottom-up mechanisms using anomaly composite analysis for SST, wind-driven Ekman upwelling, and primary production. Higher SSTs often correlate with reduced nutrient availability (Behrenfeld et al., 2006), while Ekman upwelling brings nutrient-rich waters to the surface, boosting primary productivity (P, 1995). We found different SST anomaly patterns between adults and juvenile wandering albatrosses. In adults, years with higher male survival and breeding probability for both sexes correlated with a positive SST anomaly in the central subtropical Indian Ocean and a negative anomaly off the west coast of Australia (Fig. 3B, Appendix Fig. S2,3), coinciding with stronger winds along the eastern edge of the subtropical high (Fig. 2D). No similar pattern was detected in juveniles. Both adult and juvenile years with higher vital rates (excluding juvenile male survival) showed a higher temperature gradient between the Agulhas Return Current and the subtropical fronts (Fig. 3A,B, Appendix Fig. S2-6). Years with higher adult male survival, female breeding probability, and juvenile male survival also showed positive upwelling anomalies in the zonal regions of 30-40^◦^S and 50-70^◦^S, and negative anomalies around 40-50^◦^S (Fig. 3C, Appendix Fig. S3, 4). These patterns are consistent with the upwelling anomaly composites linked to an intensified and poleward-shifted MH (Appendix Figs. S7C and S8C). For both juveniles and adults, years with higher vital rates featured positive anomalies in primary production in the region between the Agulhas Return Current and subtropical fronts (Fig. 3A,D, Appendix Fig. S2-6).

**Figure 3.**
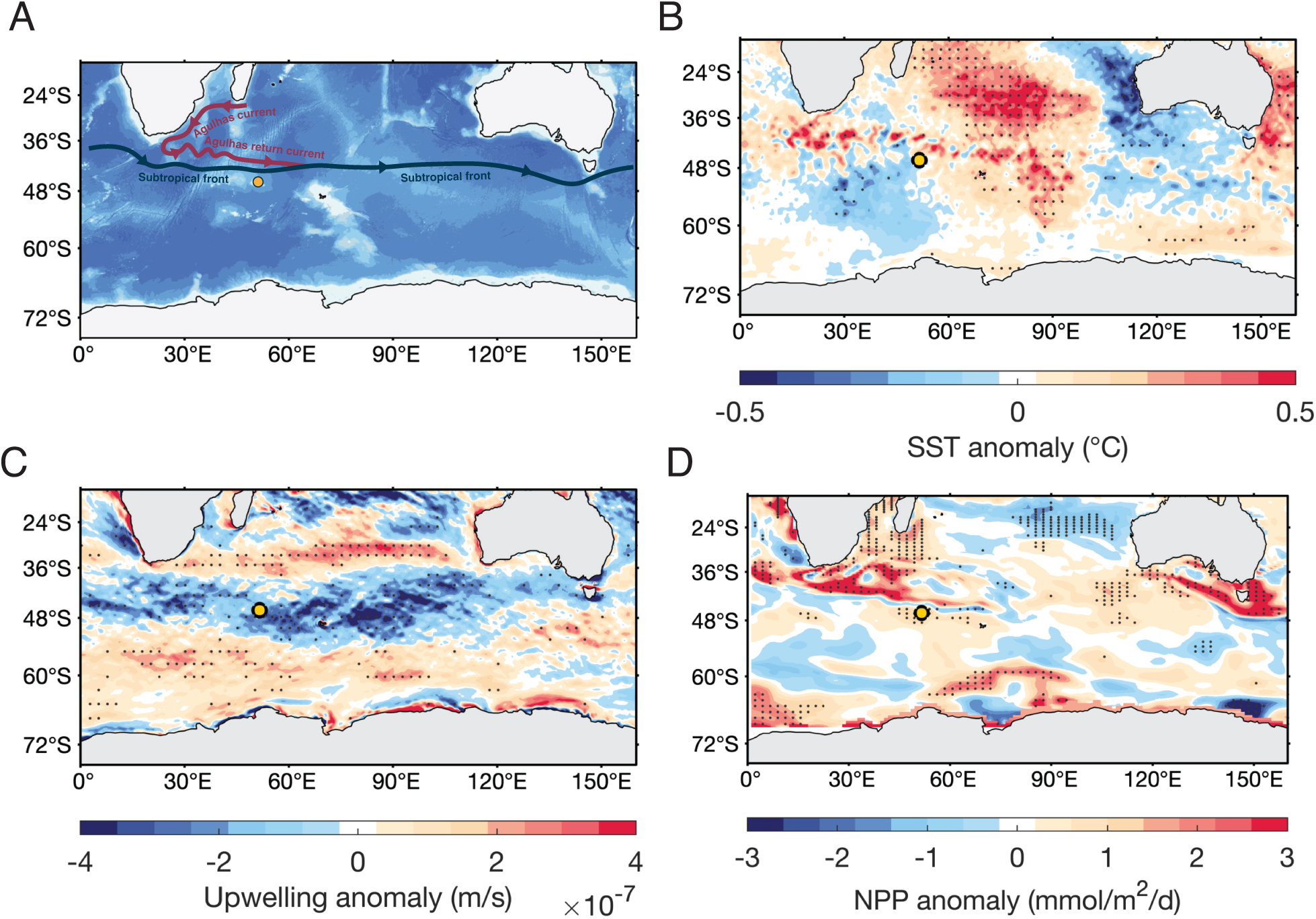
Anomaly composite analysis of potential bottom-up mechanisms influencing male wandering albatross survival probability. (A) Location of key oceanographic features, such as the Agulhas Return Current and Subtropical Front. (B), (C) and (D) show differences in sea surface temperature (SST), Ekman upwelling, and net primary productivity (NPP), respectively, between composites of high (higher 25th percentile) and low (lower 25th percentile) male survival. NPP was measured as the vertical integral of total carbon fixation between 0 and 100 m below the surface. Light dotted stippling indicates significant differences between years of higher and lower male survival probability.

## 4 Discussion

Subtropical anticyclones influence large-scale wind and weather conditions, with significant implications for ecosystems. Here, we show that variability in the location and intensity of the Indian Ocean subtropical anticyclone, the Mascarene High (MH), influences vital rates across the entire life cycle of wandering albatrosses breeding on a sub-Antarctic island south of the MH. Specifically, an intensified, poleward, and eastward-shifted MH enhances westerly winds (Fig. 4), creating conditions favorable for dynamic soaring (Richardson et al., 2018), and shaping demographic outcomes in juveniles and adults, with differences between sexes. These results underscore the importance of large-scale atmospheric systems in structuring marine predator populations and highlight subtropical anticyclones as potential drivers for understanding life history and population responses to climate variability.

**Figure 4.**
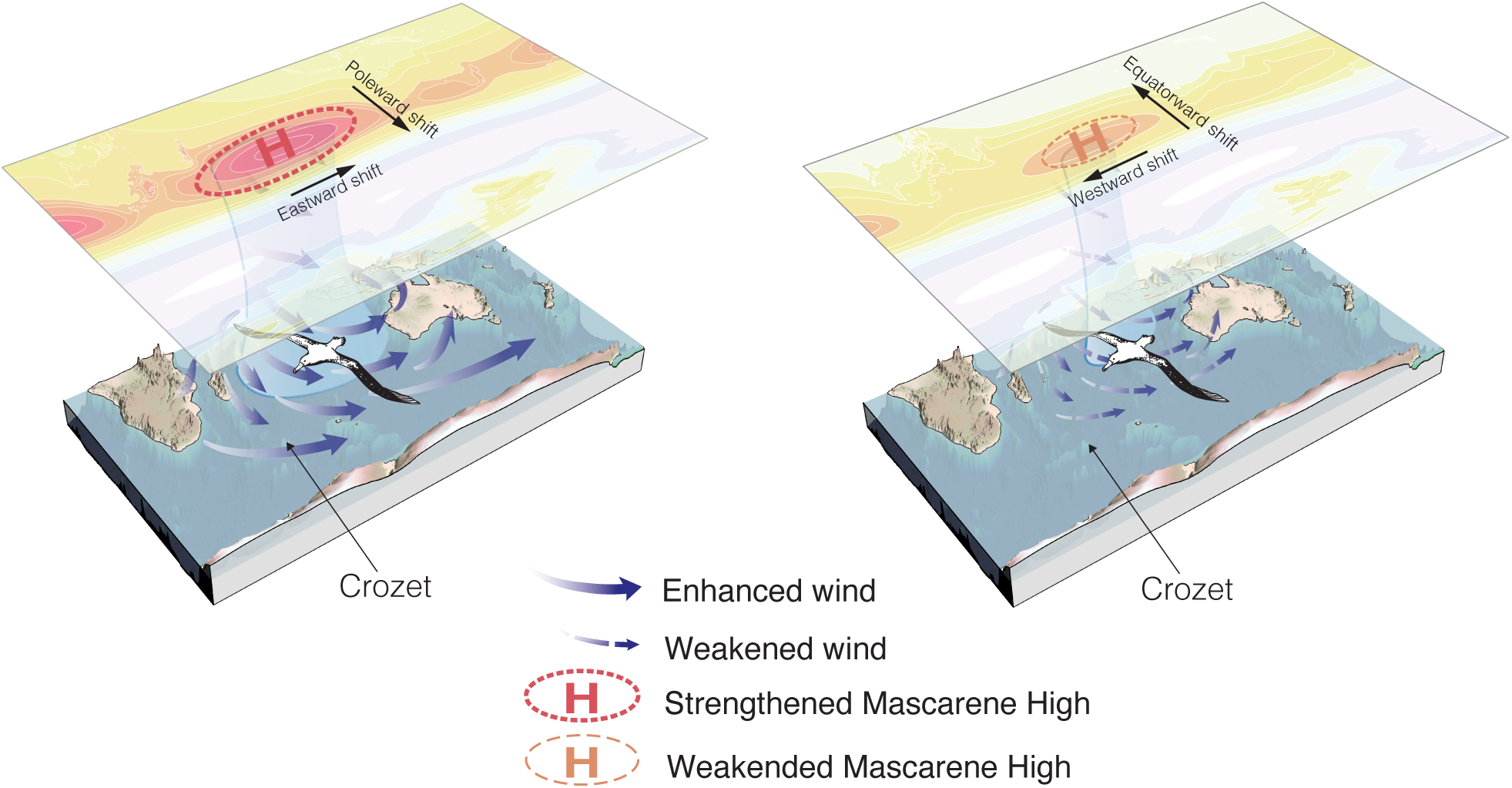
Schematic illustrating how changes in the intensity and position of the Mascarene High affect wind conditions for wandering albatrosses breeding on Crozet. A stronger, poleward-shifted Mascarene High enhances westerly winds, creating favorable conditions that improve albatross survival and reproduction.

Wandering albatrosses rely heavily on wind energy for dynamic and slope soaring (Richardson, 2011). This population of wandering albatrosses forages south of the MH, within the westerlies, with females and juveniles ranging slightly farther north than males (Weimerskirch et al., 2014). Both an intensified and poleward-shifted MH manifest as an enhanced meridional pressure gradient, compressing and poleward displacing the westerly wind belt and enhancing overall wind conditions across the southern Indian Ocean (Fig. 4). Changes in the meridional component of the subtropical anticyclone center have been shown to affect mid-latitude westerly windds in particular (Rykaczewski et al., 2015). An eastward shift of the MH moves the high-pressure center further from Crozet, enhancing winds near the colony.

The MH variability may also influence the life-history traits through bottom-up mechanisms. A stronger and poleward-shifted MH is associated with stronger meridional pressure gradients between the tropics and subtropics, promoting enhanced large-scale circulation and eddy propagation. These intensified westerly winds, along with increased trade winds, are likely to increase wind-stress curl across the Indian Ocean. Furthermore, intensified westerly winds can enhance Ekman upwelling and deepen mixed layer depth. As the western boundary current of the southwest Indian Ocean, the Agulhas Current also transports nutrients from the low-latitude tropics to the nutrient-limited subtropical and subpolar gyres in subsurface “nutrient streams” (Pelegrí & Csanady, 1991). The enhanced induction of these nutrients into the mixed layer of the gyres fuels primary production in the western part of the southern Indian Ocean. Additionally, variability in the MH affects rainfall and wind patterns over southern and eastern Africa (Manatsa et al., 2014). This can influence the transport of dust aerosols containing essential nutrients (e.g., iron), deposited into nutrient-limited ocean surfaces and boosting primary production (Gittings et al., 2024). Although we lack direct observations of biomass fluctuations in the cephalopod species targeted by wandering albatrosses, cephalopods have fast life-history traits and are known to respond rapidly to oceanographic variability and shifts in primary production (Arkhipkin, 2004; Hoving et al., 2013). Even faster responses may be expected in the small-bodied squid species most frequently consumed by albatrosses. Our use of MH indices averaged over different timescales (12 months for survival, 10 months for breeding success, and 4 months for breeding probability) likely captures these short-term changes in prey availability. Future fieldwork incorporating observations of prey species abundance will be essential to fully understand the community responses to MH variability. Future fieldwork incorporating observations of prey abundance will be essential to fully establish how MH variability propagates through the food web to shape top predator demography.

The MH effects on juveniles and adults were different, reflecting differences in life-histories. While MH variability primarily influences breeding probability in adults recovering from prior reproductive efforts, it affects breeding success in first-time breeders due to their inexperience. Previously bred adults must recover from their prior breeding attempt, which is highly reliant on environmental conditions (Weimerskirch, 2018). Thus, MH primarily affects adults by influencing breeding decisions, as experienced breeders are better equipped to handle harsh conditions during the breeding season. For first-time breeders, the initiation of their first breeding attempt relies on establishing a partnership. After fledging, they stay at sea for up to seven years before returning to their natal colony, where they begin breeding after several years of visits to bond with a partner (Weimerskirch et al., 1997). The onset of breeding is delayed until their body condition is optimal and a partnership is formed (Weimerskirch, 1992), with age being a primary determinant of probability for the first breeding attempt in juveniles (Fay et al., 2016). Once breeding begins, the MH affects first-time breeding success due to their lack of foraging efficiency, limited physiological reserves, and unfamiliarity with their partner, all of which make them more vulnerable to environmental fluctuations and less capable of successfully raising offspring (Weimerskirch, 1992).

Adult male wandering albatrosses were more affected by variability of MH than females. While MH influenced both survival and breeding probability in males, it only affected breeding probability in females, likely reflecting sex-specific differences in the benefits of certain climate conditions. Wandering albatrosses are sexually dimorphic, with males being 20% larger in mass and having a longer wingspan than females (Shaffer et al., 2001). Their greater body mass results in higher energy expenditure during takeoff, making them more dependent on strong winds (Clay et al., 2020). Additionally, adult males travel further, cover broader areas, and migrate more poleward, making them more influenced by westerly winds (Shaffer et al., 2001; Weimerskirch et al., 2012). This higher foraging effort by males may come at a survival cost, particularly in unfavorable wind conditions, and sex-specific foraging pattern responses to wind conditions have been observed across many seabird species (Lewis et al., 2015; Wakefield et al., 2009).

The MH explains the dynamics of more life-history traits in wandering albatrosses than widely used climate indices such as ENSO or IOD (Appendix Tables S1 and S2). Although MH, ENSO, and IOD are linked through complex nonlinear teleconnections (Cai et al., 2011), the MH provides a more direct and regionally relevant indicator of subtropical climate variability. Subtropical anticyclone indices have previously been shown to outperform broader climate modes in predicting regional environmental conditions. For example, the Azores High better predicts drought in the Iberian Peninsula than the NAO (Cresswell-Clay et al., 2022), and the Western Pacific Subtropical High has been shown to strongly influence the monsoon and migration patterns of the brown planthopper (Lv et al., 2023). Our findings further suggest that subtropical anticyclone indices are valuable predictors of demographic rates in wildlife populations, particularly for wide-ranging species influenced by wind-driven systems.

We demonstrated the importance of developing climate indices grounded in a thorough understanding of atmospheric and oceanographic dynamics, achieved by leveraging data and collaboration between ecology and climate sciences. A mechanistic understanding that goes beyond simple statistical correlations is essential for interpreting ecological responses, as many studies rely on correlations with climate variables without accounting for the underlying physical mechanisms driving these relationships. As highlighted in Mesquita et al. (2015), relying on simplified proxies without comprehending the underlying physical mechanisms can obscure the complex interactions between climate variability and ecological processes.

We show that variability in subtropical anticyclones effectively captures variations in wind distribution and intensity, exemplifying how a well-defined atmospheric index can reveal large-scale wind patterns and their cascading effects on ecosystems. Wind variability has recently emerged as a crucial driver of animal behavior and physiology (Thorne et al., 2023), yet quantifying its impact on fitness and population dynamics remains challenging, likely due to mismatches in spatiotemporal scales. Demographic processes such as survival and reproduction are typically assessed annually, whereas the ecological mechanisms linking wind variability to behavior and physiology operate on much shorter timescales, from hours to days (e.g., in insect (Alma et al., 2017; Hennessy et al., 2020), mammal (Renda et al., 2023; Togunov et al., 2017), and bird species (Saraux et al., 2016)). Additionally, spatial scales differ markedly. Like albatrosses, many wind-affected species, such as blue whales (Ryan et al., 2022) and common noctules (*Nyctalus noctula*) (Hurme et al., 2025), travel thousands of kilometers annually, complicating the link between local wind conditions and demographic rates. The reliance on multi-month wind averages, which correlate with large-scale annual demographic rates, may obscure underlying ecological mechanisms. Indices derived from subtropical anticyclones can help bridge this gap by capturing the dominant drivers of these relationships, filtering out noise from short-term variability, and providing a clearer understanding of how wind patterns influence population dynamics.

Moreover, the subtropical anticyclone indices link atmospheric circulation with surface wind dynamics and bottom-up processes, which are critical in many wind-driven systems. In the California Current System, influenced by the North Pacific High, blue whales track intensified wind-driven upwelling that leads to more aggregated krill population (Ryan et al., 2022). Incorporating North Pacific High indices may improve the predictability of blue whale presence and inform strategies to mitigate collision risks. Similarly, the Azores High influences Atlantic trade winds (Cresswell-Clay et al., 2022), which in turn regulate prey availability for Eleonora’s falcons (*Falco eleonorae*) and affect their breeding success (Gangoso et al., 2020). Additionally, subtropical anticyclones were also found to be a great predictor for the frequency of storms (Wang et al., 2013). Given that many pelagic seabird species respond to storms (Weimerskirch & Prudor, 2019), subtropical anticyclone indices may also serve as a valuable tool for predicting storm-driven population dynamics in these species.

Each of the five subtropical anticyclone systems in the world’s oceans regulates the physical environment of distinct and expansive ecosystems. Observations and climate models indicate that these systems have strengthened and shifted poleward over the 20th century, a trend projected to continue in the 21st century (Backeberg et al., 2012; Bograd et al., 2023; Cresswell-Clay et al., 2022; Li et al., 2013). Given the robustness of the signal in both past and future projections, incorporating subtropical anticyclone systems into ecological models can enhance ecological forecasts, supporting ecosystem management and conservation under climate change. The MH indices exemplify how integrating climate variability across temporal and spatial scales can elucidate the mechanisms linking atmospheric processes to demography. By reducing noise in species’ responses to environmental changes, the index enhances predictability and provides a versatile tool for ecological research and conservation. Looking ahead, extending the application of subtropical anticyclone indices to other wind-driven systems will enable more robust projections of ecological responses under climate change, advancing both scientific understanding and conservation.

## 5 Ethics

Licences and permissions for capture and handling of animals were granted by the Ethic Committee of Institut Polaire Francais (IPEV) and by the Préfet of Terres Australes et Antarctiques Francaises (TAAF) after advice from the Comité de l’Environnement Polaire (CEP).

## Acknowledgments

We acknowledge all the field workers involved in long-term demographic studies since 1960 on Possession Island for their invaluable help with data collection and Dominique Joubert for help with data management, as part of Project 109, “Seabirds and marine mammals as sentinels of global changes in the Southern Ocean: eco-evolutionary trends and processes,” funded by the French Polar Institute Paul-É mile Victor (IPEV; PI C. Barbraud). We acknowledge IPEV, Terres Australes et Antarctiques Françaises, and Zone Atelier Antarctique et Terres Australes (CNRS) for logistical and financial support. This study is part of the long-term Studies in Ecology and Evolution (SEE-Life) program of the CNRS and a contribution to Project SENSEI (Sentinels of the sea ice) funded by Fondation BNP Paribas. RS thanks Joanie Van de Walle and Francesco Ventura for thoughtful discussions and comments.

## Funding

National Science Foundation grant NSF-GEO/NERC 1951500 (SJ, RS, and ER);

National Science Foundation grant NSF-ORCC 2222057 (SJ, RS, and ER);

The Woods Hole Oceanographic Institution through the Joint Initiative Awards Fund from the Andrew W. Mellon Foundation (SJ and CCU);

The James E. and Barbara V. Moltz Fellowship for Climate-Related Research (CCU);

The WHOI Independent Research and Development Program (to CCU);

National Aeronautics and Space Administration (NASA) grant 80NSSC20K1289 (KK).

