## Appendix S1 for "Subtropical anticyclones shape life-history traits of a wind-reliant marine top predator"

### **Manuscript title:** Subtropical anticyclones shape life-history traits of a wind-reliant marine top predator

Ruijiao Sun[1,2,4*], Etienne Rouby[2], Christophe Barbraud[5], Henri Weimerskirch[5], Karine Delord[5], Kristen Krumhardt[6], Caroline C. Ummenhofer[3†], Stéphanie Jenouvrier[2†]

1. Marine Science Institute, University of California Santa Barbara, Santa Barbara, CA, USA
2. Biology Department, Woods Hole Oceanographic Institution, Woods Hole, MA, USA
3. Department of Physical Oceanography, Woods Hole Oceanographic Institution, Woods Hole, MA, USA
4. Department of Earth, Atmospheric and Planetary Science, Massachusetts Institute of Technology, Cambridge, MA, USA
5. Centre d'Etudes Biologiques de Chizé, CNRS-La Rochelle University UMR7372, 79360 Villiers en Bois, France
6. Climate and Global Dynamics, NSF National Center for Atmospheric Research (NCAR), Boulder, Colorado, USA

**Corresponding Author:** Ruijiao Sun

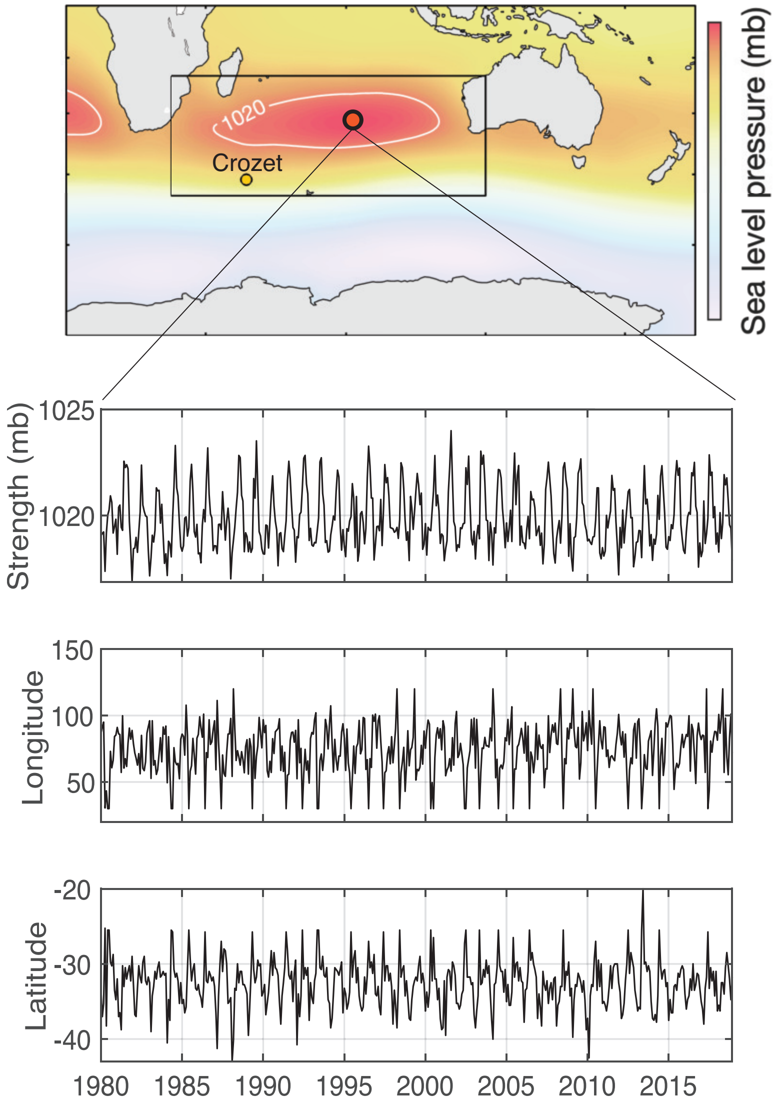


Figure S1: Definition and time series of Mascarene High (MH) indices with monthly values from 1980 to 2018. The MH area was defined as the total area within the region 30°E–120°E and 20°S–50°S (black box) above a sea level pressure of 1020.0 millibars (white contour). The center of the MH is identified as the location with the highest sea level pressure within the MH area. The MH strength refers to the average sea level pressure value of this area, and the latitude and longitude of the MH center represent the spatial location of MH.

Table S1: Effects of Mascarene High (MH) conditions on survival, breeding, and breeding success probability for adult females and males. 'Slope' indicates the effect size of MH indices at logit scale and 'f' signifies f-value.

| Demographic rate | Predictor | Female | | Male | |
|---|---|---|---|---|---|
| | | Slope | f value | Slope | f value |
| Survival probability | IOD | 0.043 | 67.4% | **0.174** | **98.5%** |
| | ENSO | -0.0415 | 70.9% | **0.113** | **94.8%** |
| | MH strength | 0.077 | 82.5% | **0.128** | **95.7%** |
| | MH latitude | -0.089 | 87.2% | **-0.132** | **97.0%** |
| | MH longitude | -0.034 | 66.0% | 0.087 | 86.5% |
| Breeding probability | IOD | -0.008 | 50.6% | -0.036 | 74.7% |
| | ENSO | -0.047 | 75.6% | -0.049 | 82.3% |
| | MH strength | 0.029 | 65.3% | **0.105** | **97.5%** |
| | MH latitude | **-0.131** | **98.3%** | **-0.102** | **98.1%** |
| | MH longitude | 0.052 | 77.1% | 0.008 | 55.5% |
| Breeding success | IOD | 0.040 | 80.3% | 0.051 | 86.9% |
| | ENSO | -0.017 | 64.2% | -0.052 | 88.0% |
| | MH strength | 0.007 | 54.7% | -0.013 | 61.1% |
| | MH latitude | -0.015 | 64.4% | -0.017 | 66.8% |
| | MH longitude | 0.011 | 59.4% | 0.043 | 84.6% |

Table S2: Effects of Mascarene High (MH) conditions on survival, breeding, and first breeding success probability for juvenile females and males. 'Slope' indicates the effect size of MH indices at logit scale and 'f' signifies f-value.

| Demographic rate | Predictor | Juvenile female | | Juvenile male | |
|---|---|---|---|---|---|
| | | Slope | f value | Slope | f value |
| Survival probability | IOD | **-0.227** | **94.6%** | -0.052 | 64.9% |
| | ENSO | -0.022 | 57.9% | 0.002 | 50.5% |
| | MH strength | -0.049 | 65.5% | -0.164 | 87.9% |
| | MH latitude | 0.018 | 57.3% | -0.103 | 75.3% |
| | MH longitude | 0.112 | 81.5% | **0.205** | **95.7%** |
| Breeding probability | IOD | 0.080 | 87.0% | -0.032 | 68.1% |
| | ENSO | 0.051 | 77.1% | -0.031 | 66.8% |
| | MH strength | -0.070 | 84.8% | -0.037 | 72.4% |
| | MH latitude | 0.022 | 62.4% | 0.015 | 59.2% |
| | MH longitude | 0.052 | 76.8% | -0.030 | 66.8% |
| | IOD | 0.010 | 54.6% | 0.025 | 63.5% |
| | ENSO | -0.098 | 86.5% | -0.084 | 87.4% |
| Breeding success | MH strength | **0.116** | **90.8%** | 0.097 | 89.1% |
| Breeding success | MH latitude | -0.016 | 58.0% | **-0.102** | **92.7%** |
| | MH longitude | 0.063 | 75.5% | **0.135** | **96.6%** |

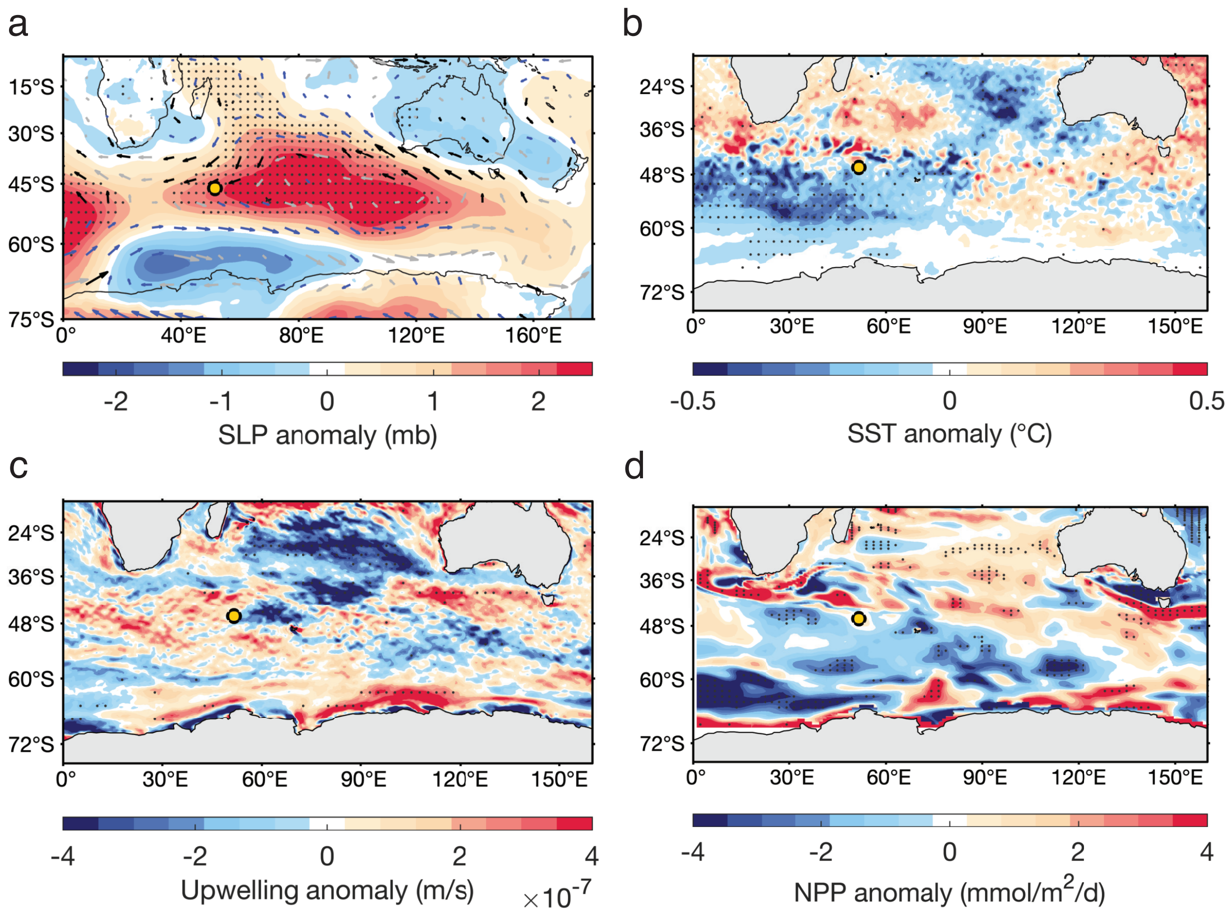


Figure S2: Composite anomaly analysis for the breeding probability of adult male wandering albatrosses. (a) Differences in sea-level pressure and wind conditions during composites of high and low male breeding probability years (25th percentile). Black wind vectors indicate significant wind differences associated with reduced wind speed, blue wind vectors indicate significant differences with increased wind speed, and light gray wind vectors indicate insignificant differences. Light dotted stippling denotes significant differences in sea-level pressure. Differences in (b) sea surface temperature (SST), (c) Ekman upwelling, and (d) net primary productivity (NPP) during composites of high and low breeding probability years of adult male wandering albatrosses (25th percentile). NPP was measured as the vertical integral of total carbon fixation between 0 and 100 m below the surface. Light dotted stippling indicates significant differences between composites of higher and lower adult male breeding probability.

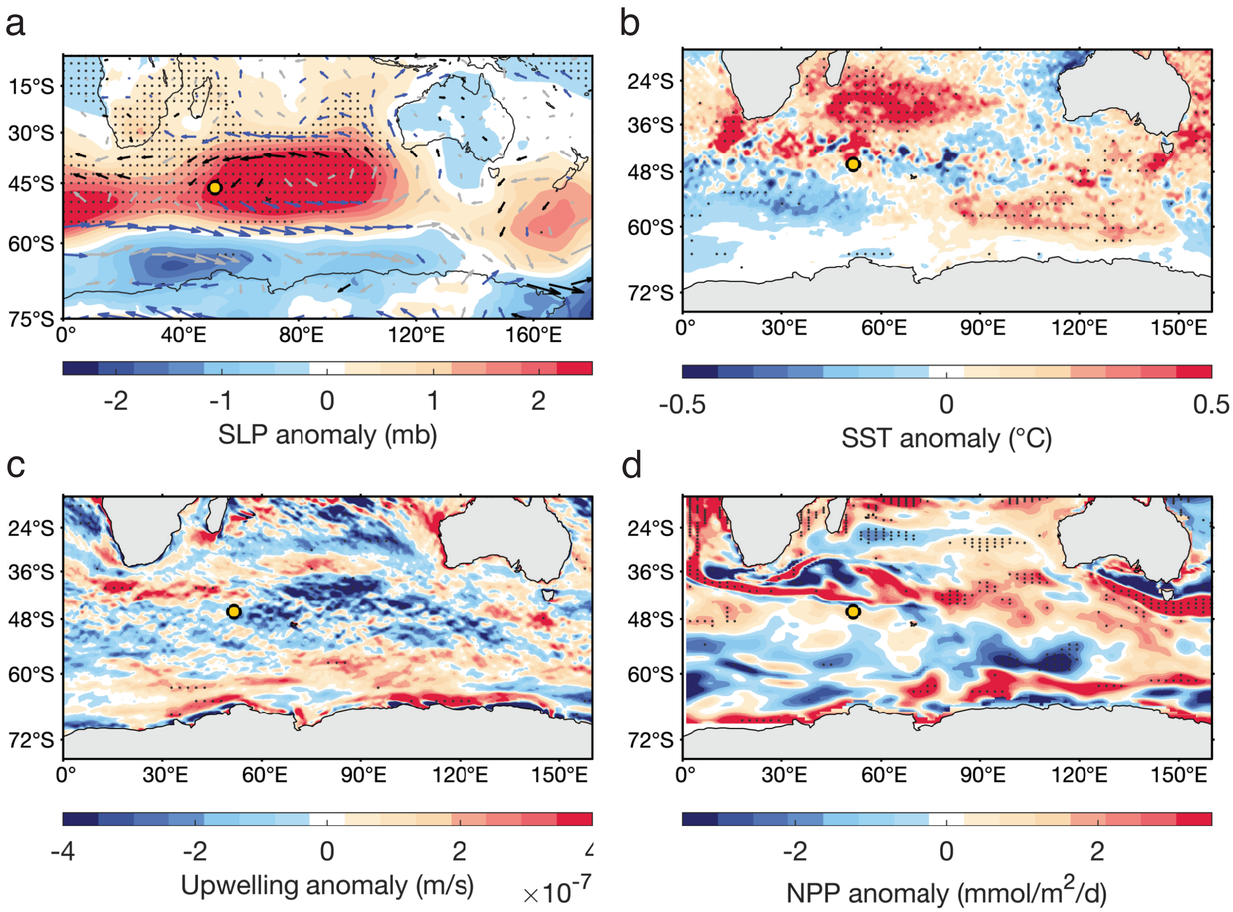


Figure S3: Composite anomaly analysis for the breeding probability of adult female wandering albatrosses. (a) Differences in sea-level pressure and wind conditions during composites of high and low adult female breeding probability (25th percentile). Black wind vectors indicate significant wind differences associated with reduced wind speed, blue wind vectors indicate significant differences with increased wind speed, and light gray wind vectors indicate insignificant differences. Light dotted stippling denotes significant differences in sea-level pressure. Differences in (b) sea surface temperature (SST), (c) Ekman upwelling, and (d) net primary productivity (NPP) during composites of high and low breeding probability years of adult female wandering albatrosses (25th percentile). NPP was measured as the vertical integral of total carbon fixation between 0 and 100 m below the surface. Light dotted stippling indicates significant differences between composites of higher and lower adult female breeding probability.

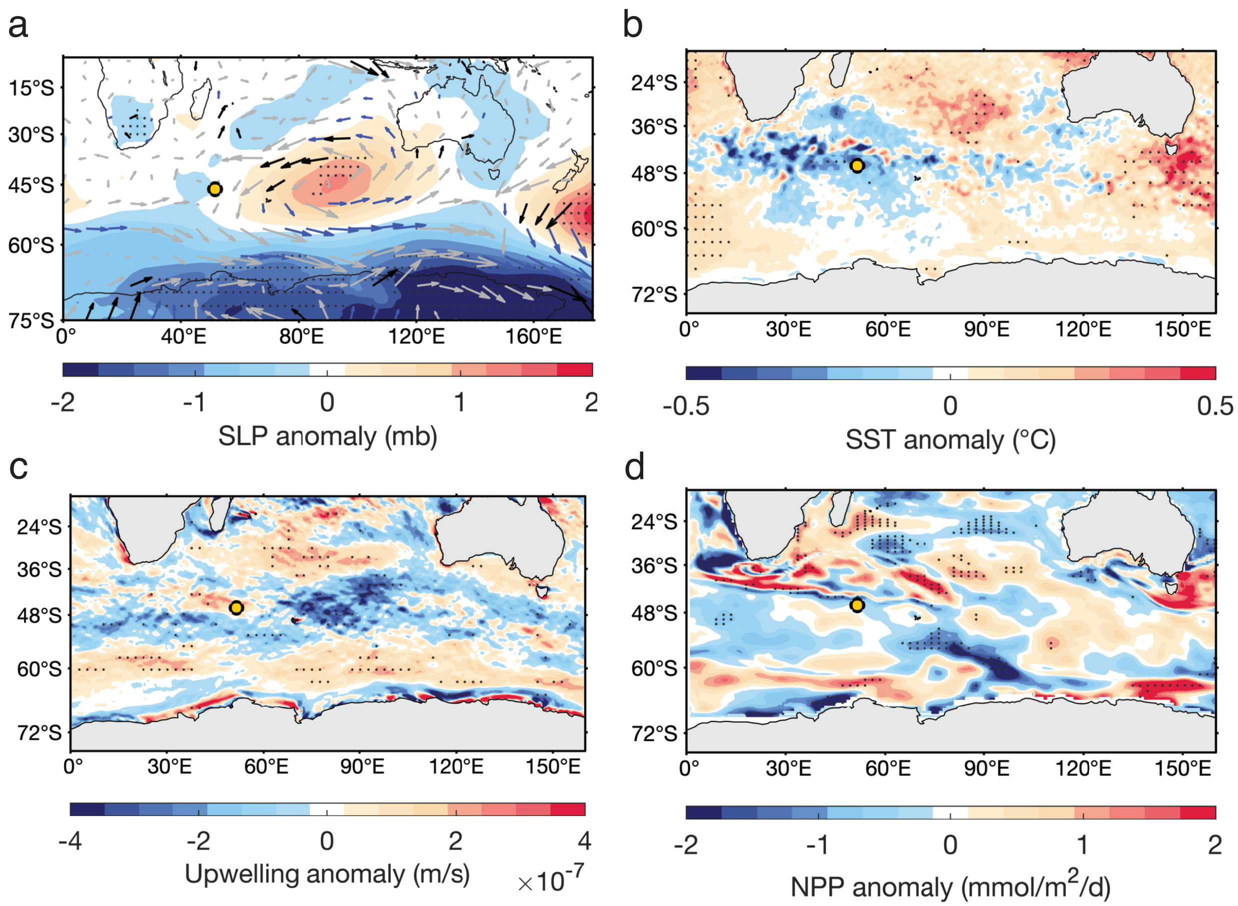


Figure S4: Composite anomaly analysis for the survival probability of juvenile male wandering albatrosses. (a) Differences in sea-level pressure and wind conditions during composites of high and low juvenile male survival probability years (25th percentile). Black wind vectors indicate significant wind differences associated with reduced wind speed, blue wind vectors indicate significant differences with increased wind speed, and light gray wind vectors indicate insignificant differences. Light dotted stippling denotes significant differences in sea-level pressure. Differences in (b) sea surface temperature (SST), (c) Ekman upwelling, and (d) net primary productivity (NPP) during composites of high and low survival probability of juvenile male wandering albatrosses (25th percentile). NPP was measured as the vertical integral of total carbon fixation between 0 and 100 m below the surface. light dotted stippling indicates significant differences between composites of higher and lower juvenile male survival probability.

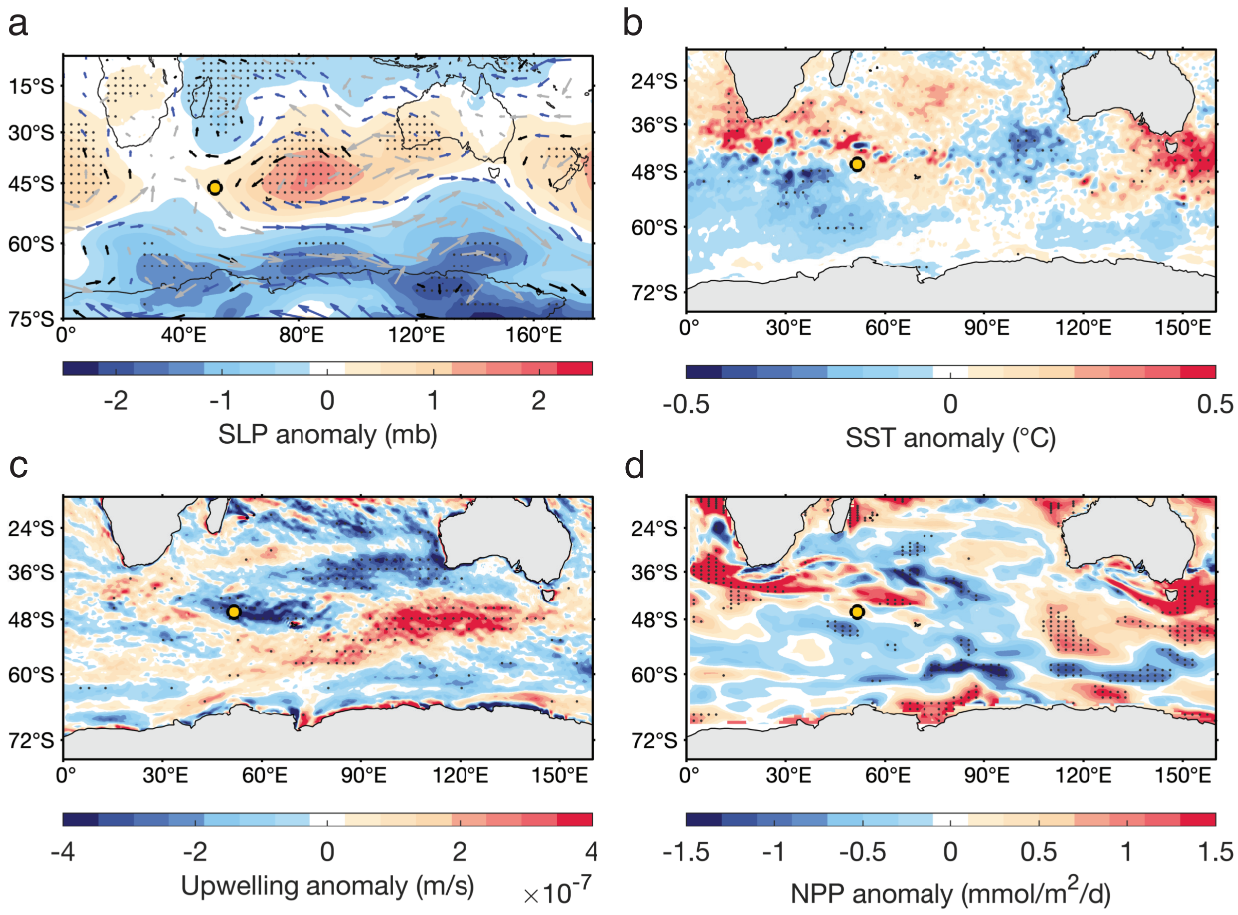


Figure S5: Composite anomaly analysis for the breeding success probability of juvenile male wandering albatrosses. (a) Differences in sea-level pressure and wind conditions during composites of high and low male breeding success probability years (25th percentile). Black wind vectors indicate significant wind differences associated with reduced wind speed, blue wind vectors indicate significant differences with increased wind speed, and light gray wind vectors indicate insignificant differences. Light dotted stippling denotes significant differences in sea-level pressure. Differences in (b) sea surface temperature (SST), (c) Ekman upwelling, and (d) net primary productivity (NPP) during composites of high and low breeding success probability years of juvenile male wandering albatrosses. NPP was measured as the vertical integral of total carbon fixation between 0 and 100 m below the surface. Light dotted stippling indicates significant differences between years of higher and lower juvenile male breeding success probability.

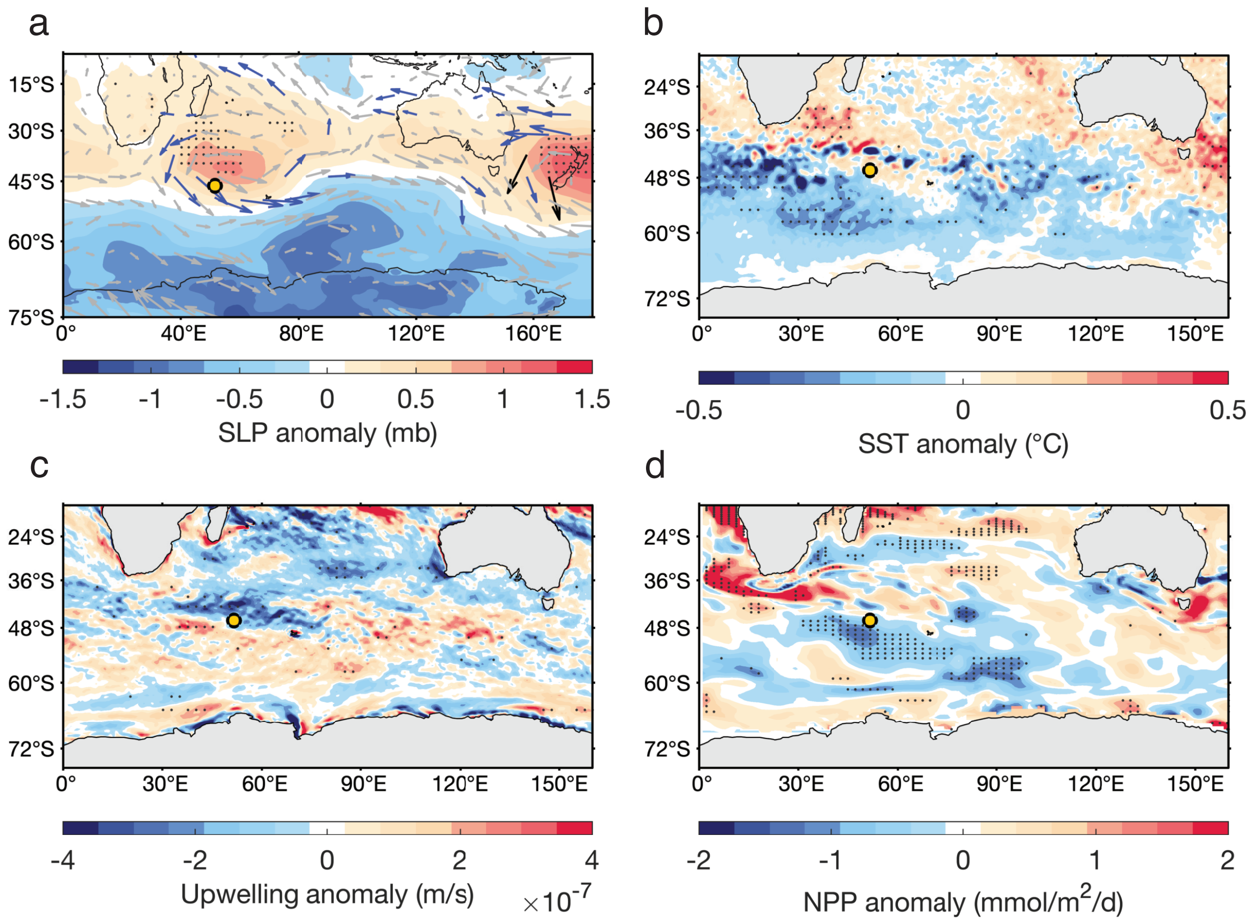


Figure S6: Composite anomaly analysis for the breeding success probability of juvenile female wandering albatrosses. (a) Differences in sea-level pressure and wind conditions during composites of high and low female breeding success probability years (25th percentile). Black wind vectors indicate significant wind differences associated with reduced wind speed, blue wind vectors indicate significant differences with increased wind speed, and light gray wind vectors indicate insignificant differences. Light dotted stippling denotes significant differences in sea-level pressure. Differences in (b) sea surface temperature (SST), (c) Ekman upwelling, and (d) net primary productivity (NPP) during composites of high and low breeding success probability composites of juvenile female wandering albatrosses (25th percentile). NPP was measured as the vertical integral of total carbon fixation between 0 and 100 m below the surface. Light dotted stippling indicates significant differences between composties of higher and lower juvenile female breeding success probability.

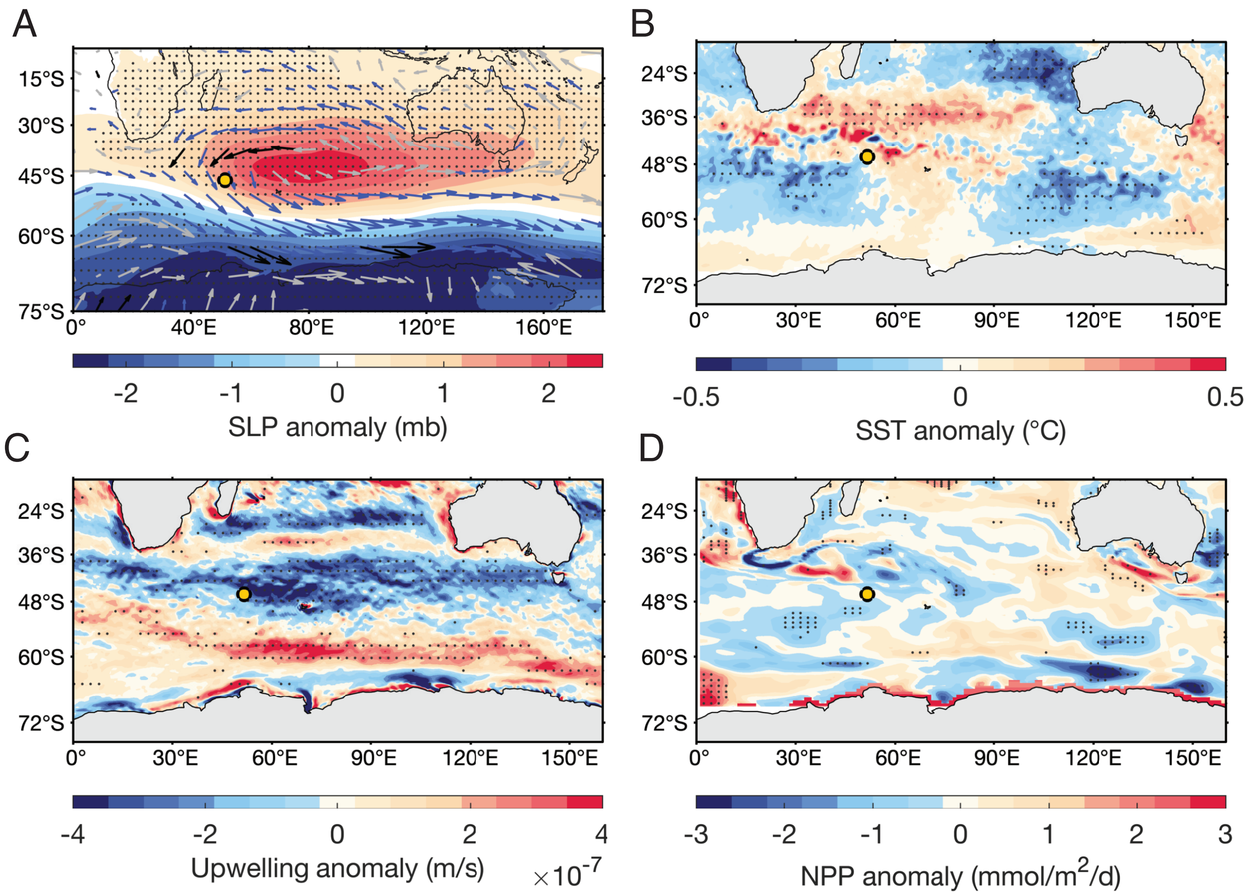


Figure S7: Composite anomaly analysis for MH strength. (a) Differences in annual mean sea-level pressure and wind conditions during composites of strong MH and weak MH (25th percentile). Black wind vectors indicate significant wind differences associated with reduced wind speed, blue wind vectors indicate significant differences with increased wind speed, and light gray wind vectors indicate insignificant differences. Light dotted stippling denotes significant differences in sea-level pressure. Differences in annual mean (b) sea surface temperature (SST), (c) Ekman upwelling, and (d) net primary productivity (NPP) during composites of strong MH and weak MH years (25th percentile). NPP was measured as the vertical integral of total carbon fixation between 0 and 100 m below the surface. Light dotted stippling indicates significant differences between composites of strong MH and weak MH.

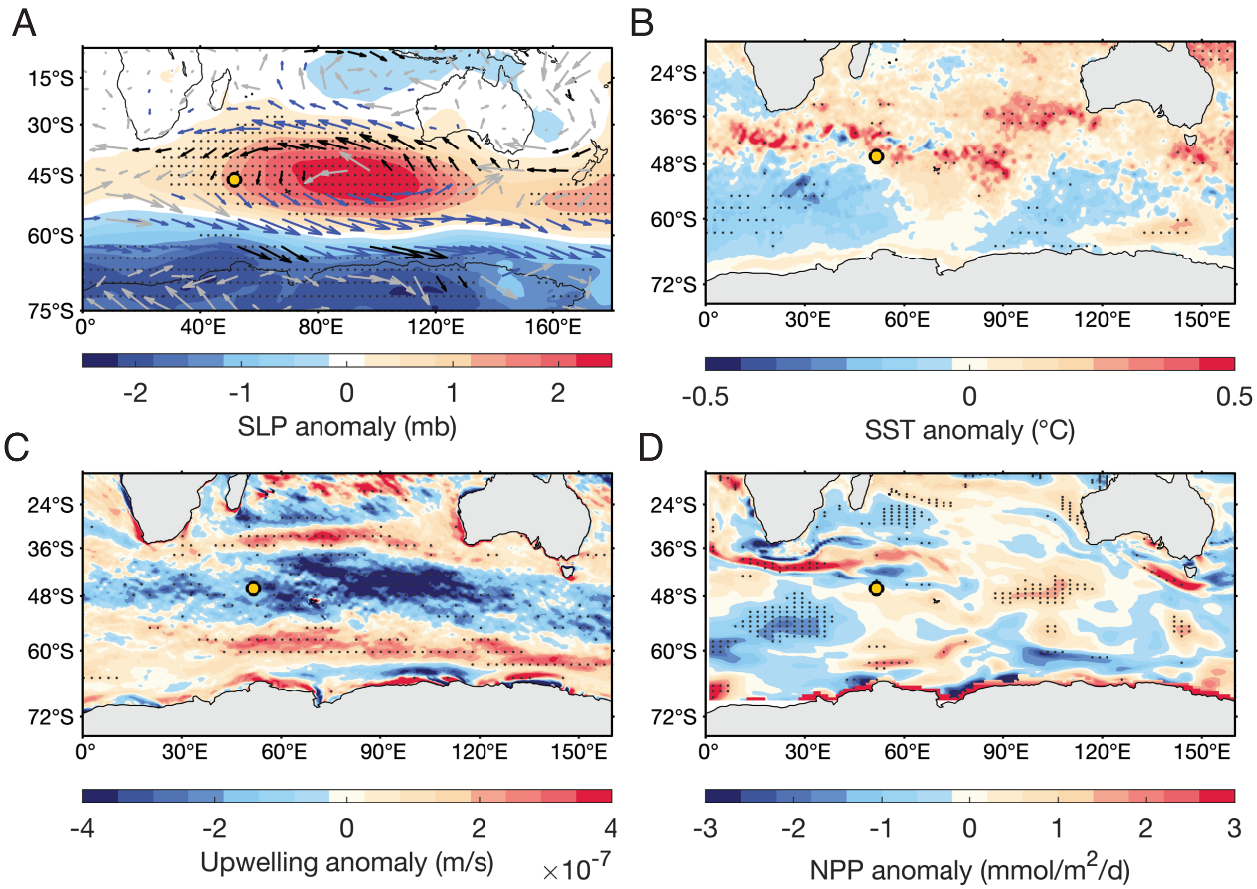


Figure S8: Composite anomaly analysis for MH latitude. (a) Differences in annual mean sea-level pressure and wind conditions during composites of poleward-shifted MH and equatorward-shifted MH (25th percentile). Black wind vectors indicate significant wind differences associated with reduced wind speed, blue wind vectors indicate significant differences with increased wind speed, and light gray wind vectors indicate insignificant differences. Light dotted stippling denotes significant differences in sea-level pressure. Differences in annual mean (b) sea surface temperature (SST), (c) Ekman upwelling, and (d) net primary productivity (NPP) during composites of poleward-shifted MH and equatorward-shifted MH (25th percentile). NPP was measured as the vertical integral of total carbon fixation between 0 and 100 m below the surface. Light dotted stippling indicates significant differences between composites of poleward-shifted MH and equatorward-shifted MH.

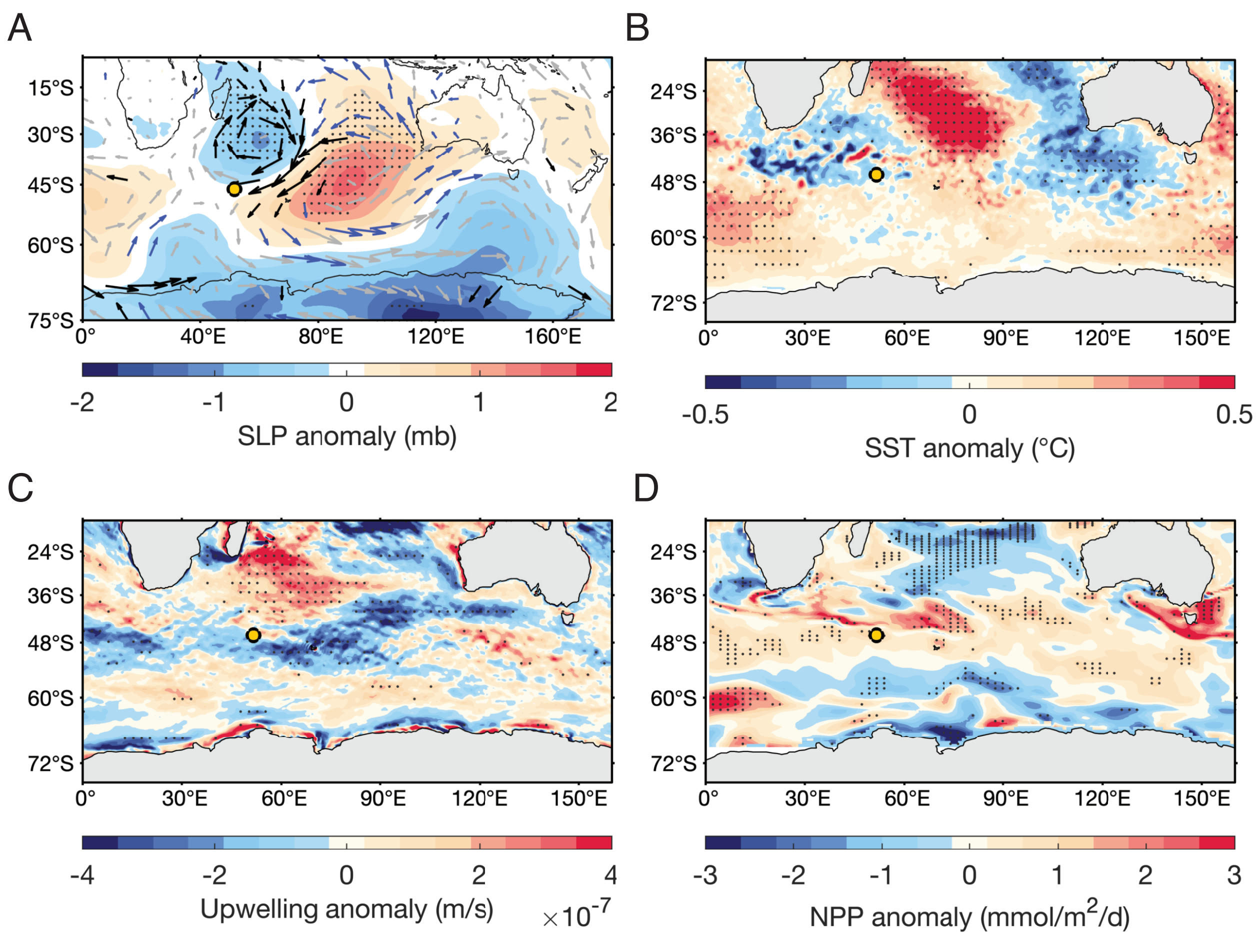


Figure S9: Composite anomaly analysis for MH longitude. (a) Differences in annual mean sea-level pressure and wind conditions during composites of eastward-shifted MH and westward-shifted MH (25th percentile). Black wind vectors indicate significant wind differences associated with reduced wind speed, blue wind vectors indicate significant differences with increased wind speed, and light gray wind vectors indicate insignificant differences. Light dotted stippling denotes significant differences in sea-level pressure. Differences in annual mean (b) sea surface temperature (SST), (c) Ekman upwelling, and (d) net primary productivity during composites of eastward-shifted MH and westward-shifted MH (25th percentile). NPP was measured as the vertical integral of total carbon fixation between 0 and 100 m below the surface. Light dotted stippling indicates significant differences between composites of eastward-shifted MH and westward-shifted MH.
